# FateNet: an integration of dynamical systems and deep learning for cell fate prediction

**DOI:** 10.1101/2024.01.16.575913

**Authors:** Mehrshad Sadria, Thomas M. Bury

## Abstract

Understanding cellular decision-making, particularly its timing and impact on the biological system such as tissue health and function, is a fundamental challenge in biology and medicine. Existing methods for inferring fate decisions and cellular state dynamics from single-cell RNA sequencing data lack precision regarding decision points and broader tissue implications. Addressing this gap, we present FateNet, a computational approach integrating dynamical systems theory and deep learning to probe the cell decision-making process using scRNA-seq data. By leveraging information about normal forms and scaling behavior near tipping points common to many dynamical systems, FateNet accurately predicts cell decision occurrence and offers qualitative insights into the new state of the biological system. Also, through in-silico perturbation experiments, FateNet identifies key genes and pathways governing the differentiation process in hematopoiesis. Validated using different scRNA-seq data, FateNet emerges as a user-friendly and valuable tool for predicting critical points in biological processes, providing insights into complex trajectories.

## Introduction

Complex dynamical systems can experience sudden shifts between states when they reach a critical threshold known as a tipping point or critical transition (1). These transitions have been extensively studied in fields such as ecology, climate science, finance, and epidemiology (2–6). Tipping points are also pertinent in the field of medicine, as seen in diseases such as diabetes and epileptic seizures (7,8). These tipping points are characterized by a sudden shift from a healthy state to a diseased condition, signaling a critical change in the system’s dynamics (9). Recognizing early warning signs of these transitions during disease progression would enable the identification of pre-disease conditions and facilitate timely medical intervention (10). Fortunately, there are universal properties of tipping points that can present themselves before a tipping point occurs (11,12). One such example is critical slowing down, characterized by a decrease in local stability and systematic changes in properties of time series data such as variance, autocorrelation and the power spectrum (13,14). These universal properties suggest the possibility for early warning signals across a wide range of scientific domains (15–18).

In developmental biology, cells undergo a variety of transitions as they differentiate. The Waddington landscape is a fundamental concept for understanding these critical transitions, as it explains how cells experience alterations in their transcriptome and epigenome while transitioning into unique cell types (19). Predicting early cell fate bias and understanding the mechanisms underlying cell decision-making are crucial for advancing cellular reprogramming (20). By deciphering the regulatory mechanisms governing cell fate decisions, we can reprogram cells into specific cell types for regenerative medicine applications, such as replacing damaged or lost cells in various tissues and organs (21,22).

In recent years, single-cell technologies have provided us with a high level of precision in studying individual cells, allowing us to observe and understand cellular changes (23). However, they only provide a snapshot of the cellular state, limiting our ability to capture the dynamic changes that occur over time (24). To overcome this limitation, computational methods based on pseudotime analysis have been developed to reconstruct the temporal progression of cells (25). These methods infer the order of cells along a trajectory based on the similarity of their gene expression. Various techniques are employed to perform pseudotime analysis, such as using the distance to the root cell or computing entropy for each cell to position them in the trajectory (26). While RNA velocity can be integrated into pseudotime analysis and provide information on the direction of cell state changes, it doesn’t always provide accurate directions due to various factors (27). Despite the promising results of all these methods, they are not yet able to detect critical transitions with high accuracy, and cannot provide specific details about the biological system changes that occur during and after the transition.

In recent years, deep learning has emerged as a powerful tool for predicting changes in complex dynamical systems (28,29). In particular, a neural network can learn to predict bifurcations by training it on a massive corpus of simulation data from dynamical systems with noise (30,31). However, these current methods cannot be applied directly to pseudotime series of scRNA-seq data for two reasons: (i) scRNA-seq data is very high-dimensional, typically on the order of thousands of genes; and (ii) pseudotime series do not contain temporal correlations typical of dynamical systems with noise, since each data point is a snapshot from a unique cell, not a single cell evolving over time.

In this study, we introduce FateNet, a novel computational model that combines the theory of dynamical systems and deep learning to predict cell fate decision-making using scRNA-seq data. By leveraging universal properties of bifurcations such as scaling behavior and normal forms (32), FateNet learns to predict and distinguish different bifurcations in pseudotime simulations of a ‘universe’ of different dynamical systems. The universality of these properties allows FateNet to generalize to high-dimensional gene regulatory network models and biological data. This approach not only provides an understanding of when cells undergo state changes but also captures the type of these transitions, identifying the characteristics of the system’s new state. By using FateNet we demonstrate how perturbing specific sets of genes can alter the type of transition a system undergoes. Notably, FateNet eliminates the need for training a model on the specific system under study and allows us to overcome the limitations of most deep learning models, which are typically restricted to the systems they were originally trained on. We test our model using simulated and biological scRNA-seq data of various sizes. Our results underscore FateNet’s ability to detect the process of cell fate decision-making, offering insights into the ongoing transitions within the system and providing information on manipulating gene sets to modify transition types.

## Results

### Bifurcation prediction in a simple gene regulatory network

To demonstrate FateNet’s accuracy, we use data generated from a simple model of a gene regulatory network (Methods). The model undergoes a fold bifurcation as the synthesis rate of the first gene increases (m_1), and a pitchfork bifurcation as the degradation rate of the genes (k_D) decreases (33). We simulate the model with additive white noise and a linearly changing parameter that leads to (i) a fold bifurcation, (ii) a pitchfork bifurcation, and (iii) no bifurcation (Fig. 1). In the bifurcation scenarios, the bifurcation is reached at pseudotime 500. At a given point in time, FateNet takes in all preceding data and assigns a probability for a fold, transcritical and pitchfork bifurcation, and a probability for no bifurcation (null). A heightening in a bifurcation probability is taken as a signal that this bifurcation is approaching. FateNet successfully signals the approach of a fold and a pitchfork bifurcation in the gene regulatory network. In the case of the null trajectory, no significant probability is assigned to any of the bifurcations.

**Figure 1:**
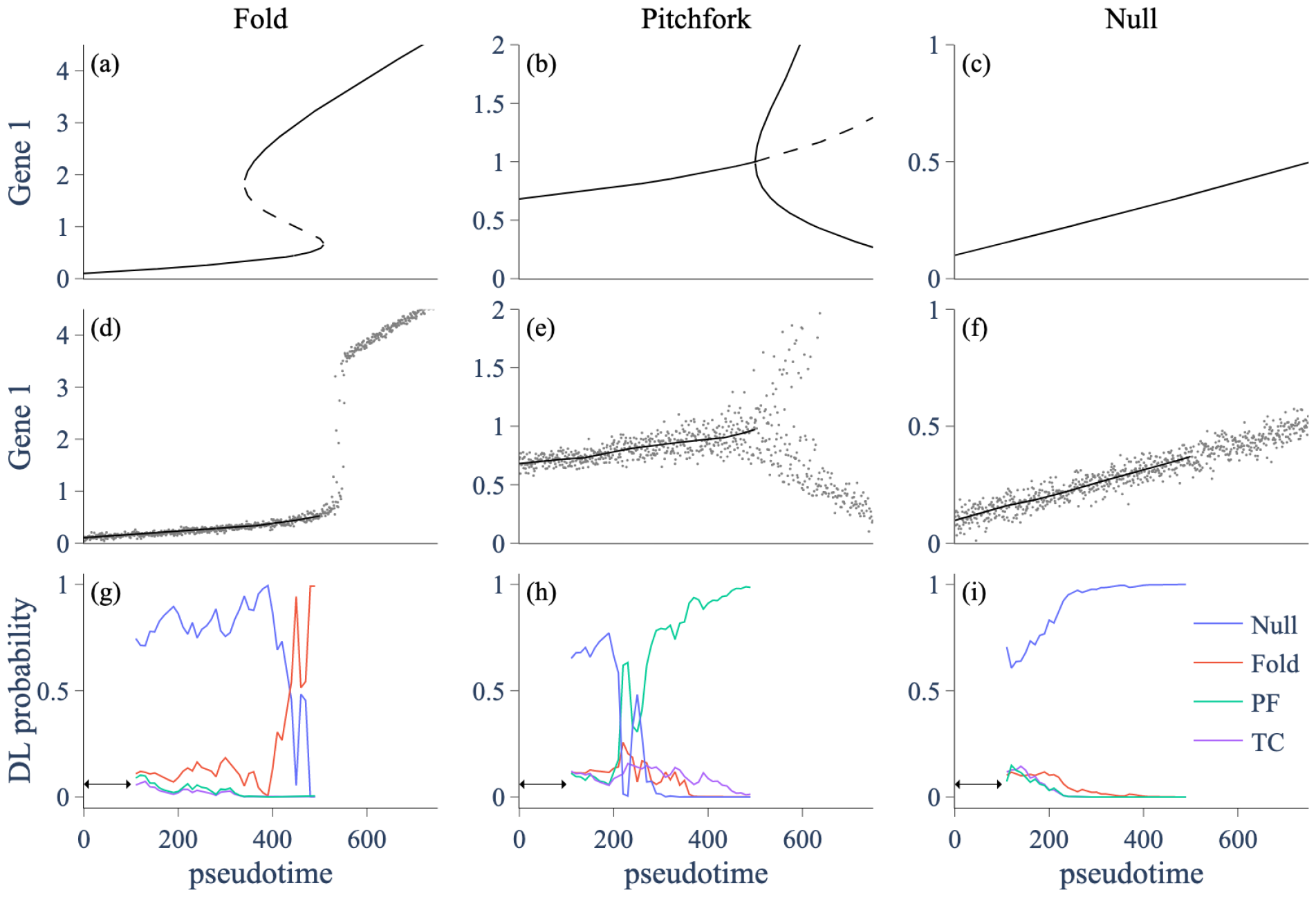
Simulations and predictions in the simple gene regulatory network model going through a fold, pitchfork, and no bifurcation. (a-c) Bifurcation diagrams showing the stable (solid) and unstable (dashed) states of the model as a parameter is varied. (d-f) Model simulation (gray) with a parameter varying according to the bifurcation diagram, and smoothing (black) with a Lowess filter with span 0.2. The model reaches the bifurcation at pseudotime 500. (g-i) Probabilities assigned by FateNet for each class of bifurcation as progressively more of the data becomes available. The algorithm only considers the data after smoothing (i.e. not the trend) when making its predictions. PF: pitchfork; TC: transcritical.

### Bifurcation prediction in a large simulated gene regulatory network

To assess the performance of FateNet on larger datasets we use SERGIO, a simulator based on stochastic differential equations that can generate both steady-state and dynamic scRNA-seq data (34). It incorporates different noisetypesfor realistic dataset creation. We generate differentiation simulation data with 1800 cells, 100 genes and 7 cell types (Fig. 2a). After the preprocessing steps in the 100-dimensional gene space, we apply Principal Component Analysis (PCA) to reduce the dimensionality of the data and obtain the principal components. To establish a cell ordering and understand the pseudotime dynamics within this dataset, we use Partition-based graph abstraction (PAGA) (Fig. 2b) (35). These data are then detrended and used as input to FateNet, leveraging the known underlying differentiation trajectory.

**Figure 2:**
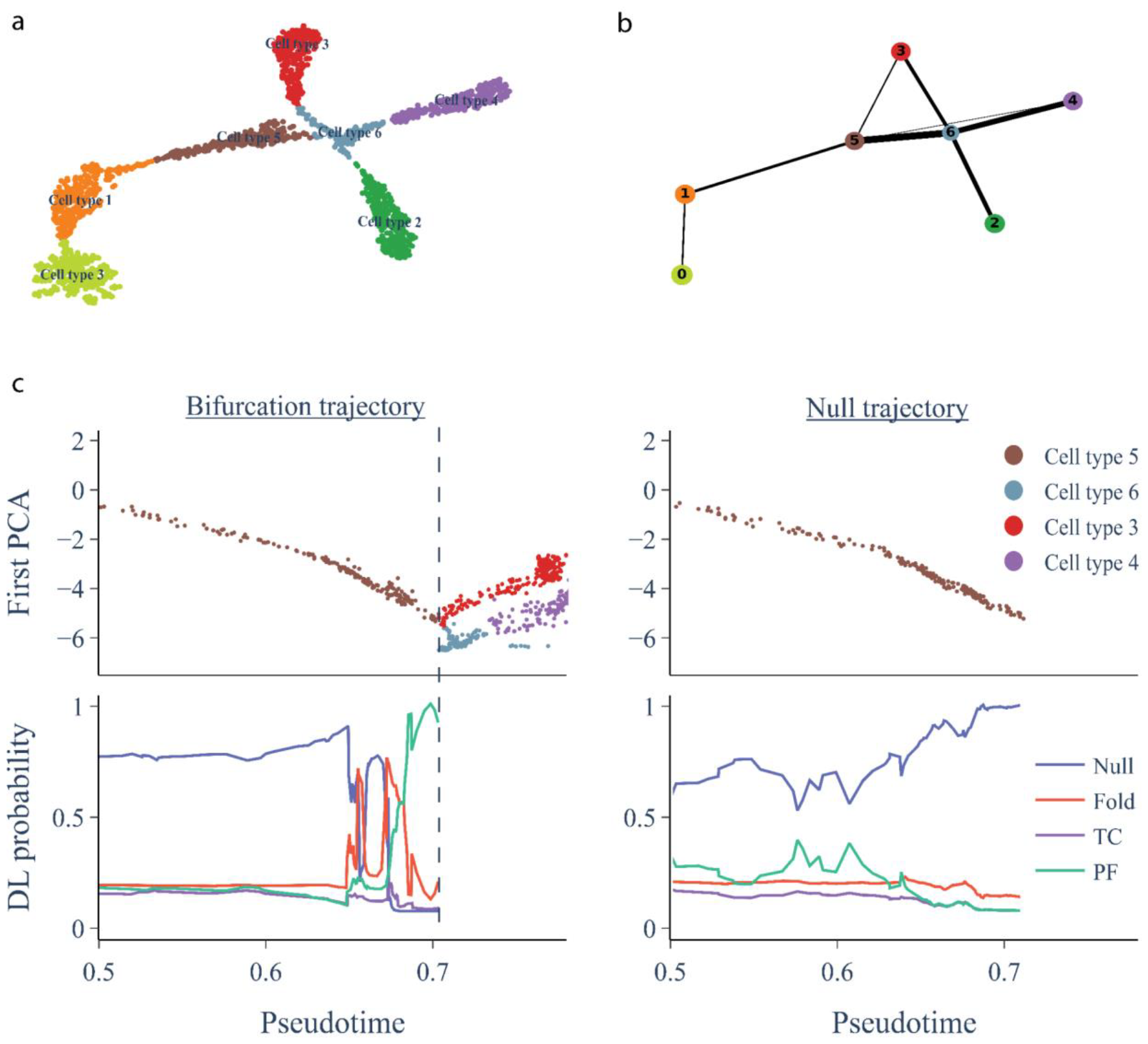
Bifurcation predictions in a simulation of SERGIO. (a) UMAP visualization of scRNA-seq data generated by SERGIO, with distinct clusters, color-coded based on cell type. (b) PAGA network graph representing the interconnectivity and relationships between cell types. (c) Bifurcation and null trajectories of cells organized in pseudotime (top) and the predictions of FateNet (bottom). The first principal component of the gene expression datais used to make predictions. The bifurcation trajectory shows a cell-fate transition between cell type 5 andcell types 3 and 6. The vertical dashedline indicates the bifurcation point. Data is smoothed using a Lowess filter with span 0.2 and the detrended data are passed to our model. The null trajectory is generated by taking a random sampling from the first 20% of the detrendeddataandaddingit to the original trend. DL probabilitiesare the probabilities assigned by our model for each event among Null, Fold, Transcritical (TC) and Pitchfork (PF).

Focusing on one of the bifurcation points, our objective is to test whether our model can predict this bifurcation in advance and identify the specific type of bifurcation occurring. We find that FateNet not only provides an early signal for the upcoming change in cell state but also successfully identifies a pitchfork bifurcation in advance, consistent with the observed change in state at the cell-fate transition (Fig. 2c, Bifurcation trajectory panel). We then test our model on a scenario where the system does not undergo a cell-fate transition (Fig. 2c, Null trajectory panel). To generate such a trajectory, we sample points randomly (with replacement) from the first 20% of the detrended data and add it to the trend of the original data. In this case, our model correctly predicts the absence of any critical transition (Null), indicating that it has learned distinct features associated with the presence/absence of an upcoming cell-fate transition (Fig. 2c, Null trajectory panel).

### Bifurcation prediction in biological data

To test our model on biological data, we use temporal scRNA-seq data of mouse hematopoietic stem cell differentiation with 130,887 cells and 25,289 genes (36). Our emphasis is on the differentiation of progenitor cells, specifically exploring the decision-making process of neutrophil fate (Fig. 3a). In this context, we aim to understand both the timing of cell fate decision-making and the specific type of differentiation occurring within the system. Therefore we use cells that are classified as undifferentiated or neutrophils and extract the top 3000 most variable genes. On this, PCA is conducted and the first principal component is used to make predictions of a bifurcation (Fig. 3b, Bifurcation trajectory panel). We find that FateNet predicts a pitchfork bifurcation before the transition from an undifferentiated cell to a neutrophil (Fig. 3b, Bifurcation trajectory panel). The transition is also preceded by an increase in variance, which is consistent with the phenomenon of critical slowing down that accompanies bifurcations. We compare this with a null time series that is generated by taking a random sample from the first 20% of the detrended bifurcation trajectory and adding this to the trend. This way, we demonstrate that the model is not making predictions based on the trend, but rather on the dynamics around the trend, which provide information about an approaching bifurcation. On the null trajectories, our model correctly predicts ‘Null’, i.e. no bifurcation (Fig. 3b, Null trajectory panel).

**Figure 3:**
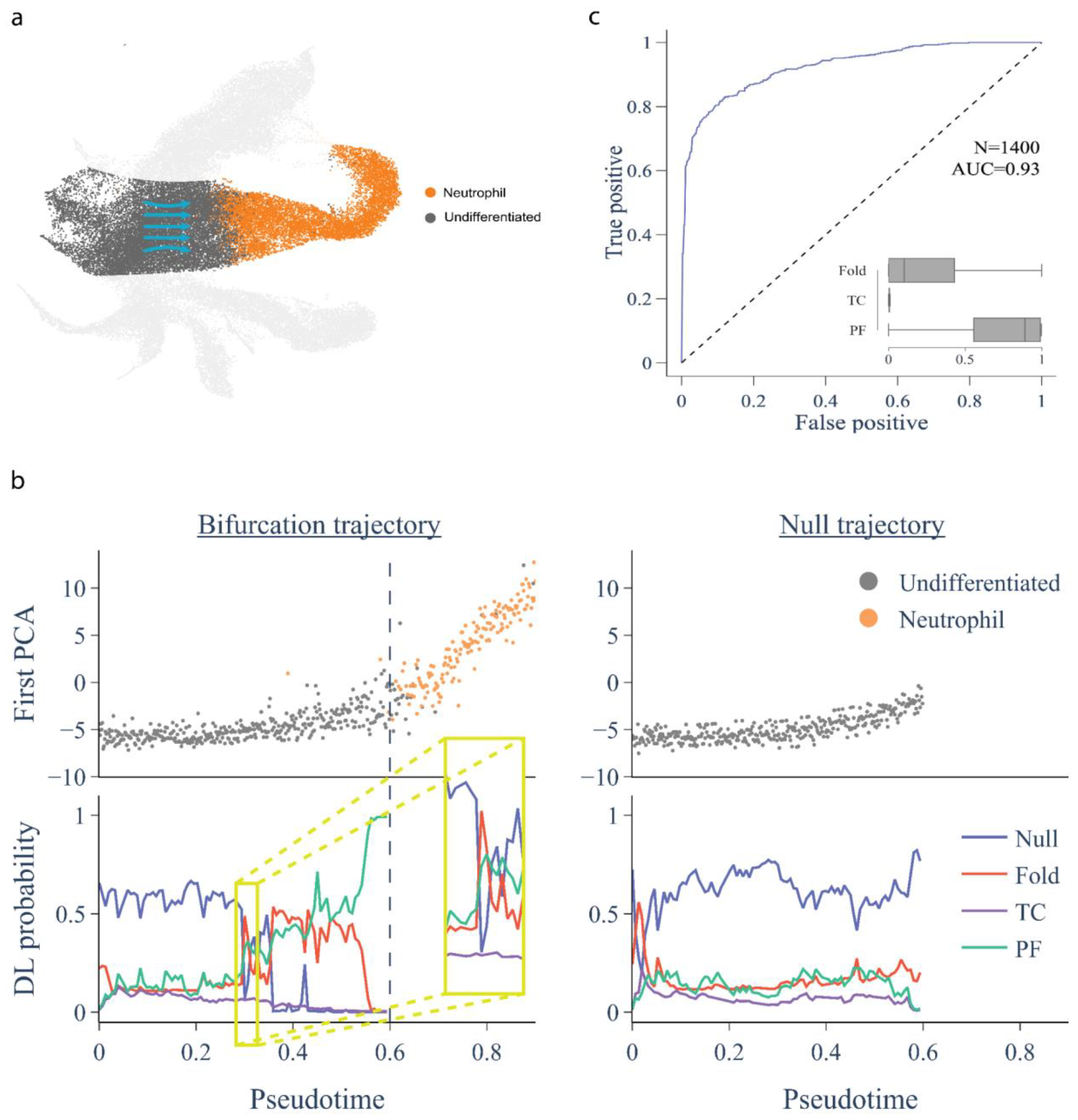
Predictions in data of mouse hematopoietic stem cell differentiation from undifferentiated cells (gray) to neutrophils (orange). (a) UMAP plot of mouse hematopoiesis data, emphasizing the transition (arrows) from progenitor cells (gray) to neutrophils (orange), elucidating the dynamic differentiation process. (b) Bifurcation and null trajectories with model predictions. The bifurcation trajectory (left) is the first principal component as a function of pseudotime downsampled by a factor of 100. The dashed line shows the transition. The data is detrended using a Lowess filter with a span 0.2 and used as input to the model. The model outputs probabilities for each event among Null, Fold, Transcritical (TC) and Pitchfork (PF). The null trajectory (right) is generated by random sampling from the first 20% of the detrended bifurcation trajectory and adding them to the trend. (c) ROC curve for 1400 evenly-spaced predictions between pseudotime 0.3 and 0.6 on 100 unique downsampled bifurcation trajectories and corresponding nulls. The inset shows the probabilities assigned to each bifurcation between pseudotime 0.5 and 0.6. Boxes show the median and interquartile range, and whiskers show the full range.

To assess the performance of our model, we make predictions on 100 unique downsampled bifurcation trajectories and corresponding nulls from the biological data. For each trajectory, we make seven equally-spaced predictionsbetween psuedotime 0.3 and 0.6, resulting in a total of 1400 predictions. The receiver operating characteristic (ROC) curve (Fig. 3c) illustrates the performance of the model at the binary classification problem of whether or not a bifurcation is approaching. An area under the curve (AUC) of 1 corresponds to a perfect performance, whereas an AUC of 0.5 (dashed line) is no better than random. In making predictions both well before and near to the bifurcation, our model achieves high performance (AUC=0.93). The prediction of the bifurcation type becomes more evident closer to the bifurcation. We show the specific bifurcation probabilities from pseudotime 0.5 onwards, demonstrating that pitchfork is the favored bifurcation across the 100 downsampled trajectories.

To understand the underlying biological mechanisms governing cell fate decision-making, we focus on a critical segment of the pseudotime trajectory specifically where a notable increase in bifurcation probability occurs (Fig. 3b, yellow box). By conducting differential gene expression analysis on cells within this specific temporal window, we identify key genes such as Myc, Ybx1, S100a8 and S100a9, Set, and H2afy whose expression showed a significant change compared to other parts of the trajectory. Remarkably, several of these genes have been shown to play a key role in fundamental cellular processes, including stem cell differentiation, regulation of neutrophil differentiation, chromatin remodeling, and cellular metabolism (Supplementary Table 1 for a detailed list of genes and their functions). Furthermore, we leverage the top 250 genes with significant expression changes, from our results to scrutinize cellular pathways, components and functions involved in cell decision-making. Our analysis reveals enrichment in pathways linked to metabolic processes (organonitrogen compound metabolism, catabolic process, superoxide anion generation, protein metabolic processes), cell death, protein localization, and leukocyte activation (Supplementary Figure 1). These findings align with existing literature showing hematopoietic stem cells navigate a complex array of developmental pathways, including not only self-renewal and differentiation but also apoptosis and metabolism. The ultimate fate of dividing stem cells is shaped by the combination of signals from various regulators. Additionally, we use the enrichment map analysis to show the network of enriched pathways, illustrating the complex relationships and communication between the identified biological processes (Supplementary Figure 2). This result not only can help us understand the connections between active pathways within the cell’s environment but also emphasizes the dynamic interactions influencing different regulatory mechanisms of cell fate decision makin g.

### Effect of gene knockout / over-expression

We investigate the effect of in silico knocking out and overexpressing genes (hard and soft interventions) on the predictions made by our model (Fig. 4a). We knockout the most significant genes in the first PCA component of the data by setting their expression to zero. We find that knocking out as few as five of the top genes results in a change in the bifurcation prediction from a pitchfork bifurcation to a fold bifurcation (Fig. 4b). Continuing to knock out genes increases the prediction for no bifurcation (Null) until eventually, after knocking out the top 30 genes, no bifurcation is predicted at all. We also overexpress genes by multiplying their expression by a factor of two. We find that overexpressing a small number of the top genes (5-10) strengthens the prediction of a pitchfork bifurcation (Fig. 4c), whereas overexpressing a larger number of genes weakens the prediction of a pitchfork bifurcation. These results suggest that there are a few genes that are instrumental in the type of bifurcation that the system goes through. When these key genes are subjected to knockout, there is a substantial alteration in the bifurcation type of the system’s dynamics (Fig. 4b). On the other hand, with the top genes overexpressed, the bifurcation type is predicted with greater probability (Fig. 4c). However, it is essential to highlight that overexpressing a broader set of genes can trigger additional regulatory mechanisms (Fig. 4a,c). This broader activation manifests in an increased probability of other types of bifurcations.

**Figure 4:**
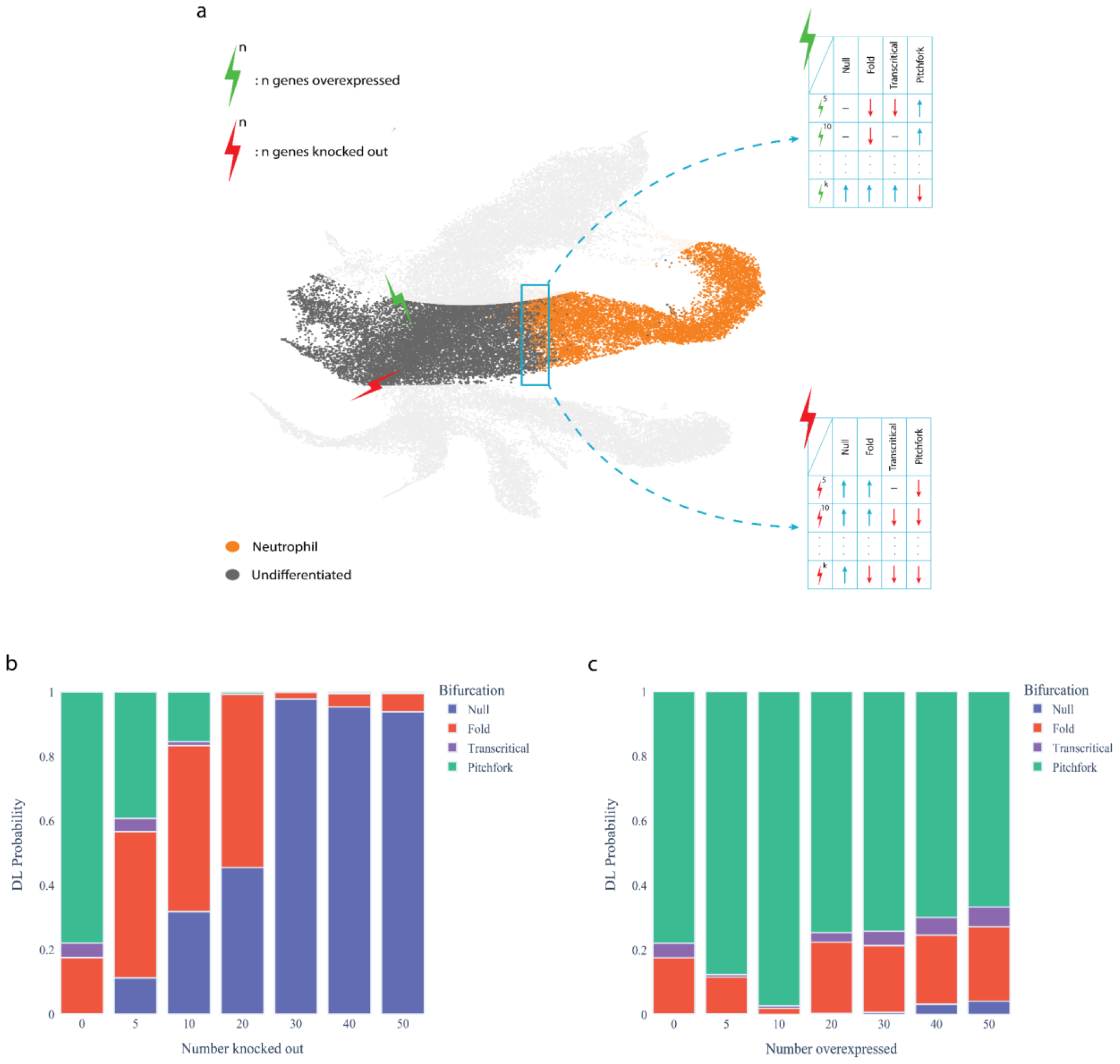
Exploring system response to various in silico perturbations. (a) UMAP visualization shows in silico perturbations, with green lightning indicating overexpression and red lightning denoting knockout perturbations. Each perturbation is individually implemented to observe how the system experiences shifts in the bifurcation dynamics. Model predictions for stem cell differentiation to neutrophils in mouse hematopoiesis after knocking out (b) and over-expressing (c) a few numbers of the most significant genes. Genes are knocked out by setting their expression to zero. Genes are overexpressed by multiplying their expression by a factor of two. In each case, ten equally-spaced predictions are made between pseudotime 0.45 and 0.6.

## Discussion

In various domains such as ecology, climate, health, and finance, the identification of critical points, often referred to as bifurcation points, holds significant importance. Early detection is crucial for strategic decision-making and intervention, minimizing the potential for adverse consequences. Also understanding the type of these transitions is pivotal for preemptive actions in these diverse fields (37). For instance, in health, the heart can spontaneously transition from a normal rhythm to a dangerous one, known as a cardiac arrhythmia. Early detection of thesecritical transitions in cardiacactivity can enable prompt medical intervention, significantly impacting patient outcomes and preventing life-threatening situations (38).

Biological processes exhibit similar critical points and can undergo different types of transitions which occur during both normal development and disease progression. In normal developmental trajectories, cells move through Waddington’s landscape, experiencing bifurcations regulated by genetic and environmental cues. These events shape cell fate, determining whether a cell adopts a neuronal, muscular, or another specialized identity (39). Similarly, in disease trajectories, bifurcations can lead to different outcomes. For instance, consider a disease process where a cell can either recover or progress to a more severe state (40). Analyzing the paths cells take during differentiation and the process of cellular decision-making, including the precise timing of these events, is crucial for understanding development and unlocking the potential of stem cell therapies (41). However, the challenges in predicting the precise type and timing of these transitions persist due to the intricate and dynamic nature of biological systems, alongside the added complexity introduced by the high-dimensional nature of the data.

In response to these challenges, we introduce FateNet, a novel framework that integrates dynamical systems theory with deep learning to discern when cells make decisions and predict the type of transition the system is approaching. For a deep learning classifier to be effective and applicable across diverse scenarios, it necessitates training on a broad spectrum of data. We generate time series training data using simulations from a comprehensive library of different dynamical system models that possess various types of bifurcation. The universal properties of bifurcations, manifested in time series as a system approaches a bifurcation, facilitate this generalizability. To validate our framework, we conduct extensive testing using both simulated and biological data, spanning different dataset sizes and varying noise levels. Since FateNet accurately predicts the type of bifurcation, it can identify genes that can prevent harmful bifurcations in a system and promote favorable transitions. Here, by performing both hard (knock-out) and soft (over-expression) interventions to a developmental system we showed it’s possible to target a specific set of genes to promote, prevent or modify the type of transition in hematopoiesis.

We believe future studies can benefit from the application of FateNet to compare bifurcation types in biological systems, particularly when comparing differences in the typeand timing of systemtransitions in healthy and diseased conditions. Many disorders, including cancer and fibrosis, can be perceived as distinct types of transitions within a biological system (42–45). Consequently, FateNet provides valuable insights into identifying effective targets for interventions and optimal timing in guiding and redirecting the system’s evolution from an impaired to a healthy state.

## Methods

### Generation of training data for FateNet

We generate training data using simulations of a library of generated dynamical systems. Each dynamical system consists of the normal form for a bifurcation and higher-order polynomial terms with random coefficients and additive white noise. We include fold, transcritical and pitchfork bifurcations in the library.

The model framework for the fold bifurcation is

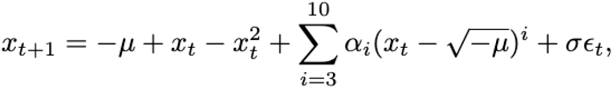

where *x*_*t*_ is the state variable, *μ*_*t*_ is the (potentially time-dependent) bifurcation parameter, *α*_*i*_ are drawn from the standard normal distribution, *σ* is the noise amplitude drawn from a uniform distribution between 0.005 and 0.015, and the noise process *ε*_*t*_ is drawn from a standard normal distribution. The initial value for the bifurcation parameter *μ*_*0*_ is drawn from a uniform distribution with lower and upper bounds that make the dominant eigenvalue of Jacobian between 0 and 0.8 (the bifurcation occurs when this eigenvalue is 1). Consequently, in the fold model, *μ*_*0*_ can take values between -0.25 and -0.01. In all models, the bifurcation occurs at *μ*=0.

The model framework for the transcritical bifurcation is

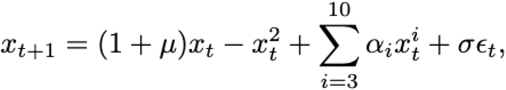

where *μ*_*0*_ can take values between -1 and -0.2. The model framework for the pitchfork bifurcation is

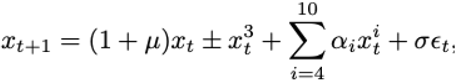

where *μ*_*0*_ can take values between -1 and -0.2. For each model framework, we generate 1000 unique models. We run 20 simulations of each model going up to its bifurcation by incrementing *μ*_*t*_ linearly from *μ*_*0*_ up to 1 over 600 time steps. If the model undergoes a noise-induced transition (defined as a deviation from equilibrium greater than 10 times the noise amplitude, *σ*), then the point prior to the transition is taken as the end of the time series. The last 500 data points are kept. If the model transitions before 500 points, it is discarded and replaced by a newly generated model. From these 20 simulations of 500 points, we construct 20 pseudotime series by placing the data in a 20x500 matrix and extracting elements in a diagonal fashion such that subsequent points in the pseudotime series come from different time simulations. Formally, denoting the data point from simulation i at time t as *x*_*i,t*_, for i=0,…,20, t=0,…,499, then the ith pseudotime series is given by *y*_*i,t*_ = *x*_(*i*+*t*)*mod20,t*_.

This process generates 20,000 ‘forced’ pseudotime series for each bifurcation. We similarly generate 20,000 ‘null’ pseudotime series where *μ* is kept fixed in each model. This gives a total of 80,000 time series that are labeled according to whether they are ‘fold’, ‘transcritical’, ‘pitchfork’ or ‘null’ trajectories. This set of time series is then shuffled and partitioned into a training, validation and test set according to the ratio 0.95:0.025:0.025. The validation and test sets were chosen as a small percentage because a set containing a few thousand time series is adequate to provide a representative estimate of the performance measures used to assess the algorithm.

### FateNet architecture and performance

FateNet is made up of two neural networks that are trained to predict a bifurcation label given a portion of pseudotime series data. Network 1 is trained on time series censored at the beginning and at the end by a randomly drawn length. This forces it to predict bifurcations based on the middle portions of the time series. Network 2 is trained on time series censored only at the beginning, allowing it to learn from data right up to the bifurcation. Formally, the length of the censored time series L is drawn from a uniform distribution with lower and upper bounds of 50 and 500, respectively. Then, for Network 1, the start time of the censored time series is drawn from a uniform distribution between 0 and 500 -L, and for Network 2 the start time is set to 500-L. The censored time series are then normalized by their mean absolute value and prepended with zeros to make them 500 points in length—a requirement for the neural network. Our model uses the average prediction from these two networks.

Each network has a CNN-LSTM (convolutional neural network—long short-term memory network) architecture (Supplementary Figure 3). This involves passing the input time series *x* ∈ *ℜ*^*500*^ through a convolutional layer to obtain the hidden units

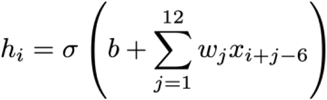

where i runs from 0 to 500, *σ* is the ReLU activation function, and b and *w*_*j*_ are the bias and weights of the kernel, which are trainable parameters. We use a kernel size of 12, and pad the edges of the time series with zeros to maintain the input dimension. We apply this operation for 50 kernel filters, resulting in a hidden layer *H*_*1*_ of dimension 500x50 and 650 trainable parameters. We apply a dropout of 10% to this hidden layer, which randomly fixes this proportion of hidden units to zero at each iteration of the training process. The units are then passed through a max pooling layer, which takes the maximum value over a window that strides over the units. We use a pool size of 2 and a stride length of 2, resulting in a second hidden layer *H*_*2*_ of dimension 250x50.

This is then passed to the first LSTM layer with 50 memory cells, where each cell is capable of capturing both long and short-term dependencies in the data (46,47). A LSTM cell consists of several components including the input gate (*i*_*t*_), the forget gate (*f*_*t*_), the output gate (*o*_*t*_), the cell state (*c*_*t*_) and the hidden state (*h*_*t*_), where *t* runs from 0 to 250. Each LSTM cell is updated as follows. The forget, input, and output gates are update as

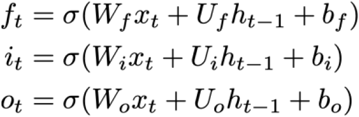

where *x*_*t*_ is the input from the previous hidden layer, W and U are weight matrices of dimension 50 x 50, b are bias vectors of length 50, *σ* is the sigmoid activation function

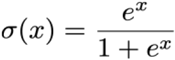

and the initial value for the hidden state is *h*_*0*_ = *0*. Meanwhile, a cell input activation vector is computed as

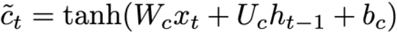

which is used to determine the new cell state and hidden state

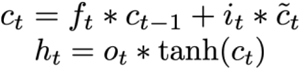

where * is element-wise multiplication of vectors. The initial value for the cell state is taken as *c*_*0*_ = *0*. The weights and biases for this layer of LSTM cells constitute 20200 trainable parameters. The resulting sequence of cell states for each memory cell is stored in the third hidden layer *H*_*3*_ of dimension 250 x 50.

This is then passed to the second LSTM layer with 10 memory cells and 2440 trainable parameters. Here, only the final value of each cell state sequence is stored, resulting in the hidden layer *H*_*4*_ of length 10. Finally, this is passed through a dense layer with softmax activation to obtain the probabilities

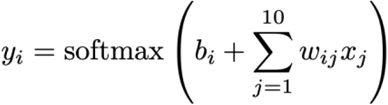

where i runs from 1 to 4, *b*_*i*_ are the biases, and *w*_*ij*_ are the weights, and *x*_*j*_ are the inputs from the previous hidden layer. These weights and biases constitute 44 trainable parameters. In total, the network contains 23,334 trainable parameters.

The networks were initialized and trained in Tensorflow v2.10 using the Adam optimization algorithm with a learning rate of 0.0005, a batch size of 1024, and categorical cross entropy as a loss function, given by

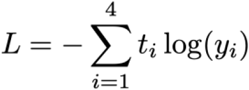

where the vector *t* is the one-hot encoded truth label and *y* is the probability vector output from the model. The network reached peak performance on the validation set after about 300 epochs of training. When evaluated on the test set, Network 1 obtained an F1 score of 0.63 on the multi-class prediction problem and 0.80 on the binary prediction problem. Network 2 obtained F1 scores of 0.92 and 0.98, respectively. The confusion matrices (Supplementary Figure 4) show the performance of the networks at predicting each class in the test set.

### Applying the model to pseudotime data

To obtain model predictions on pseudotime data, we first obtain the principal component from a PCA. We then detrend it using a Lowess filter with a span of 0.2. Predictions from our model at a given point in pseudotime are obtained by taking the preceding data, normalizing them, prepending them with zeros to make the input 500 points in length, and feeding it into the model. The model then outputs a probability vector, whose components indicate the probability of each event. This is achieved using the Python package ewstools (48).

### Simple model for a gene regulatory network

This model consists of two coupled differential equations (33) adapted to include additive Gaussian white noise and an additional parameter to modulate the nonlinear interaction between the genes. The model is given by

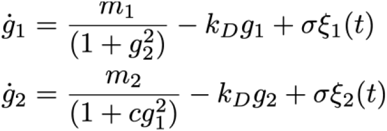

where *g*_*1*_ and *g*_*2*_ are the level of gene expression for each gene, *k*_*D*_ is the degradation rate, *m*_*1*_ and *m*_*2*_ determine the scales of their synthesis, c governs the nonlinear response of gene 2 to gene 1, *σ* is the noise amplitude, and *ξ*_*1*_ (*t*) and *ξ*_*2*_(*t*) are Gaussian white noise processes.

The model possesses a fold and a pitchfork bifurcation. We simulate trajectories going through both types of bifurcation, as well as a null trajectory where there is no bifurcation but still a time-dependent parameter. For the fold trajectory, we take *m*_*1*_ increasing linearly from 1 to 4.75, *m*_*2*_=3, *k*_*D*_=1, and c=1. For the pitchfork trajectory, we take *m*_*1*_=1, *m*_*2*_=1, *k*_*D*_ decreasing linearly from 1 to 0.25, and c=1. For the null trajectory, we take the same parameters as the fold trajectory, except we set c=0.1, which removes the fold bifurcation. In each case, we simulate the model using the Euler-Maruyama method with a step size of 0.01 and a noise amplitude *σ*=0.05. We then downsample the data by a factor of 100. As described in the section on generating training data, we then construct a pseudotime series using 20 simulations of the model and taking subsequent points from each simulation.

### SERGIO simulation

SERGIO is a simulator of single-cell gene expression data based on a stochastic differential equation (SDE) framework that can simulate noise and variability of gene expression, as well as the effects of external stimuli and cell cycle progression. SERGIO can model the stochastic nature of transcription, the regulation of genes by multiple transcription factors, and the differentiation of cells along complex trajectories. SERGIO also allows users to specify a gene regulatory network and various parameters that control the simulation process. Using SERGIO, we generated a synthetic scRNA-seq dataset that mimics a dynamic differentiation program, similar to the DS11 dataset described by (34). Our dataset consists of 1800 single cells from 7 cell types, with a gene regulatory network of 100 genes.

### Analysis of scRNA-seq data

As a preprocessing step, we scaled, centered, and log-normalized the scRNA-seq gene expression data, and extracted the top 3000 most variable genes. We used the top 40 principal components to map the highly variable genes onto a lower dimension for clustering, using a k-nearest-neighbor graph with K=30. Clusters were visualized using the Python package “UMAP”. For the scRNA-seq data, partition-based graph abstraction (PAGA) was employed to sort cells accurately over time, allowing for a detailed mapping of the dynamic progression of the processes. PAGA is an extended version of diffusion pseudotime that considers disconnected graphs. By covering both aspects of clustering and pseudotemporal ordering it can model the underlying biological progression, assigning each cell a pseudotime value that reflects its position along the inferred trajectory. Also, Cytoscape and Enrichment Map are used to investigate and visualize the connection between the identified biological pathways (49,50).

In testing our model, we consider the bifurcation from undifferentiated cells to neutrophils. The bifurcation is crossed at approximately pseudotime 0.6. For making predictions, we consider the first PCA component as a function of pseudotime up to the bifurcation. This still contains a lot of data (61309 cells), so we downsample by a factor of 100 to obtain a shorter time series more appropriate for our model. We then detrend the data using a Lowess filter with a span of 0.2. We are able to obtain 100 unique bifurcation trajectories from the biological data by shifting the downsampling procedure one point at a time. We construct 100 null time series (that do not undergo a bifurcation) by sampling randomly from the first 20% of the detrended data and adding it to the trend of the original data.

To investigate the effect of gene knockout, we set the expression of a fixed number of genes to zero. The genes that are selected are those that are most highly represented in the top PCA component. Once these gene expressions have been set to zero, we recompute the top PCA component to obtain a new bifurcation trajectory. To investigate the effect of gene overexpression, we follow a similar procedure, except we multiply the expression of the most significant genes by a factor of two.

## Data availability

The simulated scRNA-seq data can be generated by SERGIO and can be found at: https://github.com/PayamDiba/SERGIO. The hematopoiesis Weinreb et al. data can be downloaded from Gene Expression Omnibus (GEO) under accession number GSE140802. The simulated scRNA-seq data generated by the simple gene regulatory network can be found on our page.

## Code availability

Our Python implementation can be found at:

