## Supplementary material for "FateNet: an integration of dynamical systems and deep learning for cell fate prediction": Supp file

### Supplementary Information

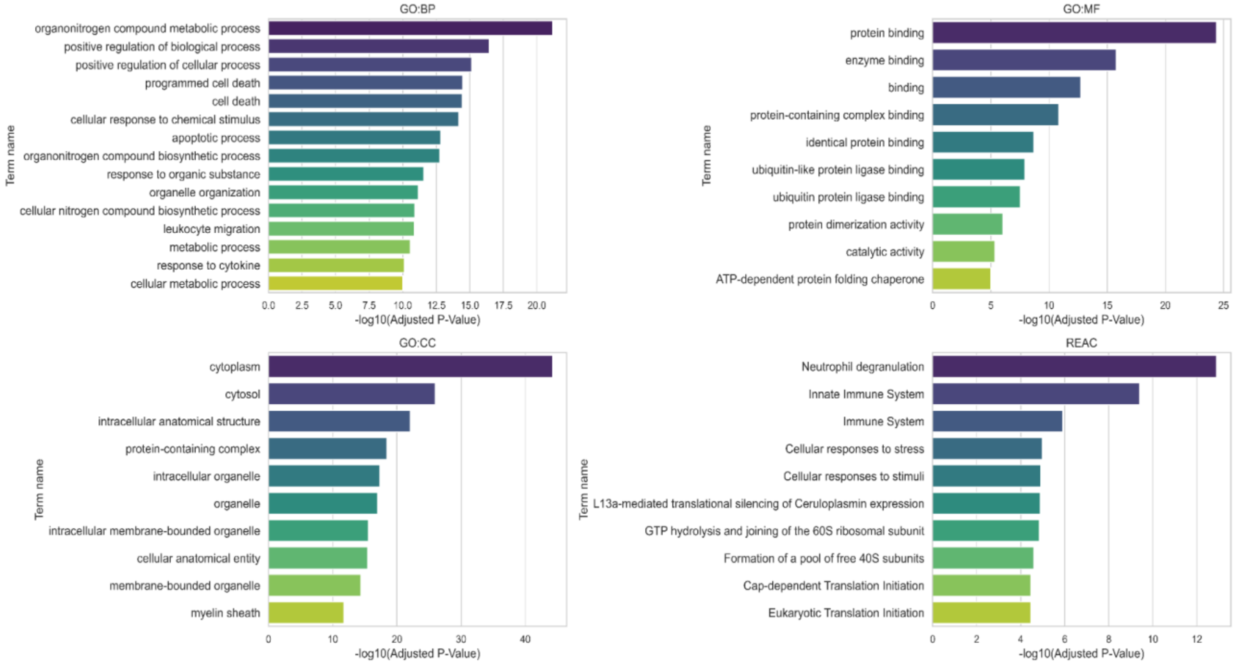


*Supplementary Figure 1. Gene Ontology (GO) term and Reactome pathway enrichment analysis for the top 250 significant genes. a) GO: Biological Process (BP) showing the enriched biological processes such as different metabolic processes, cellular response to different stimulus, and cell apoptosis indicating relevant biological activities associated with these genes. b) GO: Molecular Function (MF) highlights the enriched molecular functions including protein binding and enzyme binding. c) GO: Cellular Component (CC) displays the enriched cellular components such as cytoplasm and intracellular organelles, suggesting the subcellular localizations where these genes are active. d) The REAC panel represents the enriched pathways from the Reactome database, including immune system processes and cellular responses to stress, which are related to the process of neutrophil development.*

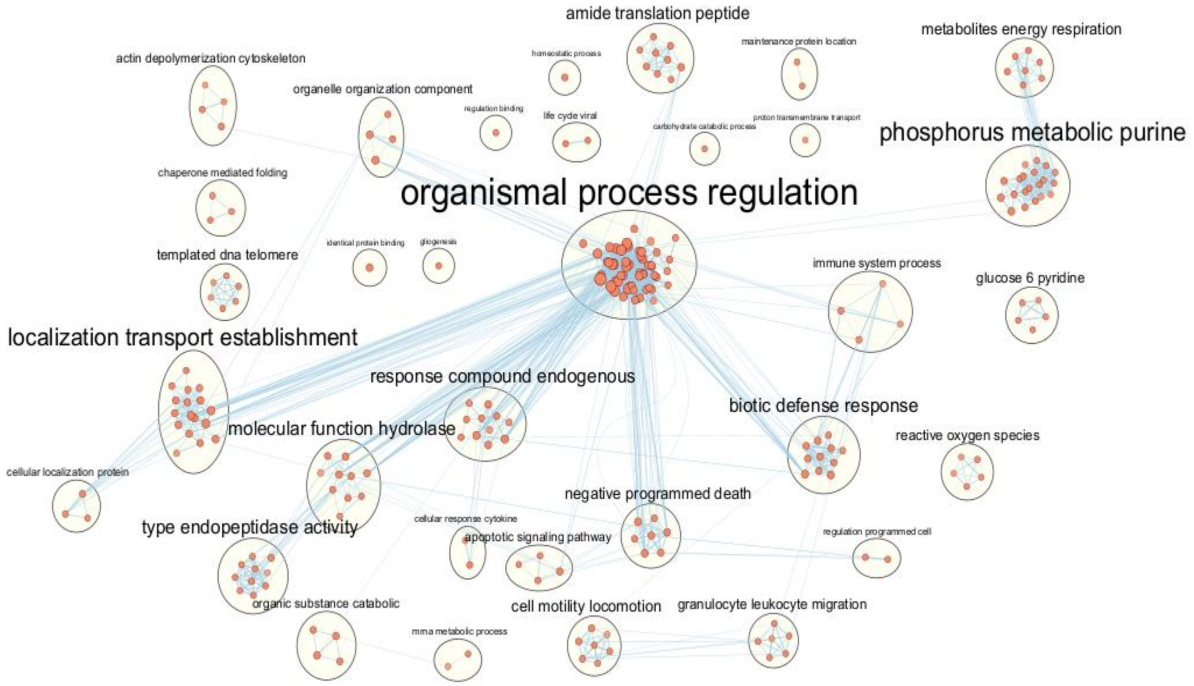


*Supplementary Figure 2. Enrichment map analysis of biological pathways. The figure shows the complex connections and interactions between different active biological processes. Blue edges of varying thickness connect the nodes, indicating the strength of the connection between pathways by the number of mutual genes. To simplify the network, only edges and nodes meeting the cutoff (Node Cutoff P-value > 0.01, Edge Cutoff > 0.35) are shown.*

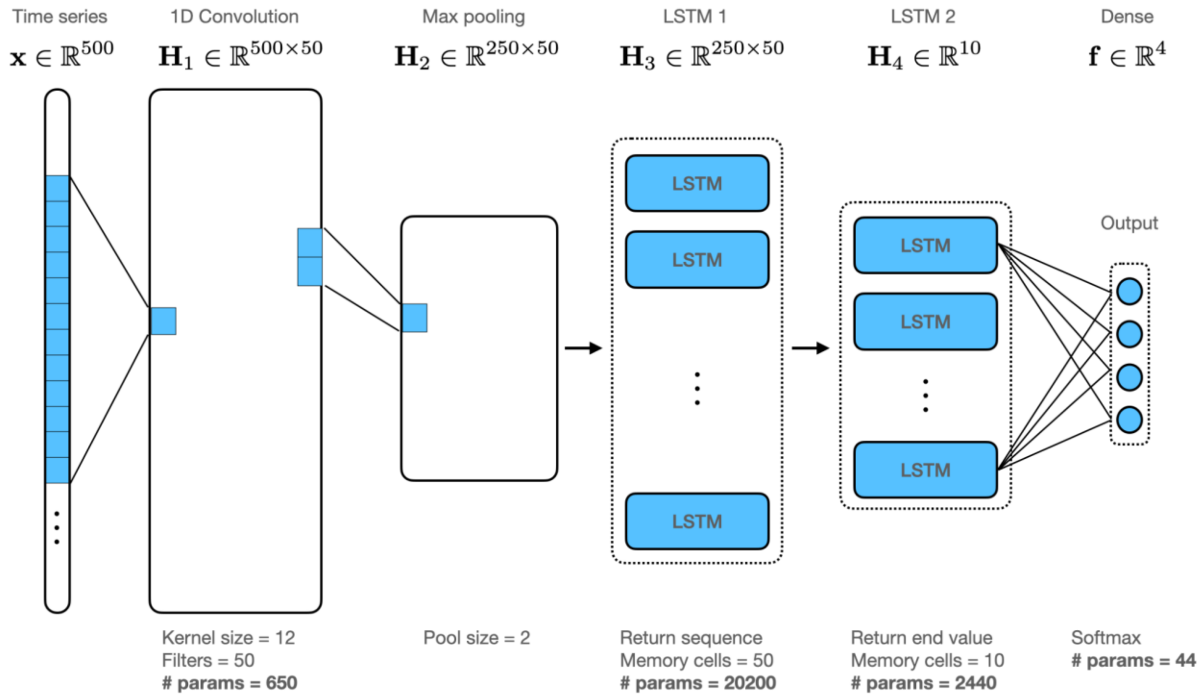


*Supplementary Figure 3. Neural network architecture of FateNet. Each neural network consists of convolutional and long-short-term memory (LSTM) layers. The input to the network is a time series of length 500. This is passed through a 1D convolution, with a kernel size of 12, 50 filters, and the ReLU activation function. Padding is applied to the ends of the time series to maintain the input dimension. This is then passed through a max pooling operation with pool size 2, reducing the dimension by a factor of 2. This is then passed to the first LSTM layer with 50 memory cells. For this layer, the cells return the entire sequence. This then enters the second LSTM layer with 10 memory cells, which returns only the end value of the sequence. Each memory cell is then connected with the 4 output nodes via a dense layer. The output is passed through a softmax filter to obtain a probability distribution over the 4 possible outcomes. The total number of trainable parameters in the network is 23,334.*

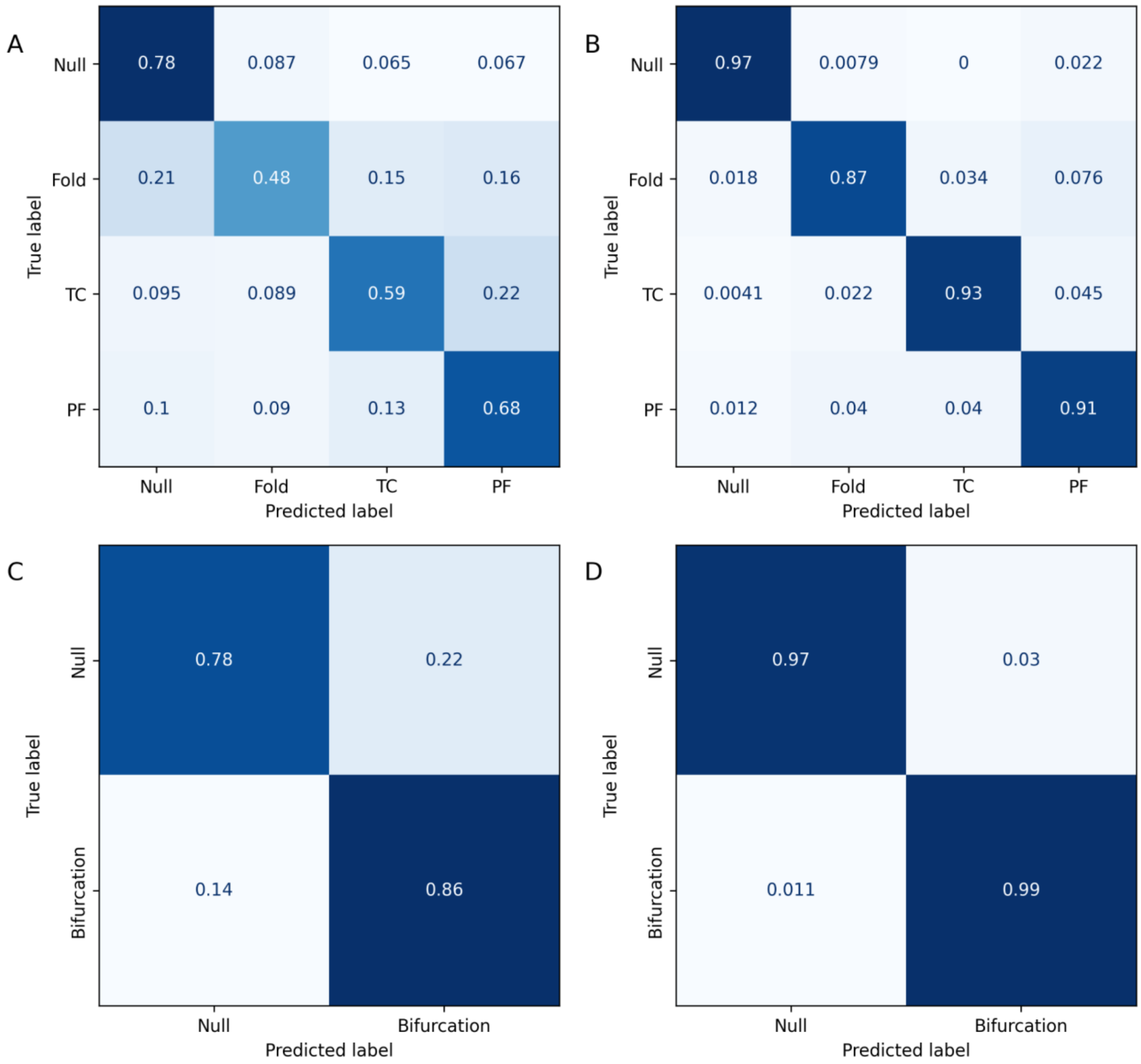


*Supplementary Figure 4. Confusion matrices for Network 1 and Network 2 when evaluated on their test sets from the library of generated dynamical systems. We show performance on the multi-class classification problem, where the model must predict the specific event, and the binary classification problem, where the model only needs to predict whether or not a bifurcation is approaching. Cell values show (row-)normalized classification rates for each class. (A) Network 1 on the multi-class classification problem obtains an F1 score of 0.63. (B) Network 2 on the multi-class classification problem obtains an F1 score of 0.92. (C) Network 1 on the binary classification problem obtains an F1 score of 0.80. (D) Network 2 on the binary classification problem obtains an F1 score of 0.98. TC: transcritical. PF: pitchfork.*

| Gene name | Reference | Role |
|---|---|---|
| Ybx1 | PMID: 21369783 | Ybx1 is a transcription factor that plays a key role in directing the commitment and differentiation of hematopoietic stem cells |
| Tmsb4x | PMID: 33310787 | Tmsb4x depletion hinders cell migration and differentiation and interferes with coronary vessel development |
| Set | PMID: 33296674 | Embryonic Stem Cell Differentiation is Regulated by SET through Interactions with p53 and β-Catenin |
| Myc | PMID: 31951039 | Myc is a family of regulator genes that code for Tfs. They are indispensable in the generation of granulocyte/monocyte progenitors (GMPs) from HPSCs |
| H2afy | PMID:21969577<br>PMID: 16428466 | It is involved in chromatin remodelling and gene expression regulation. Also, modulate transcription factor binding, X-chromosome inactivation, and transcription repression. |
| S100a9 | PMID: 28137827 | Encodes a calcium-binding protein, important in immune responses, particularly in myeloid cells. S100A8/A9 induces neutrophil activation, including adhesion, CD11b upregulation, and shedding of CD62L. |
| Prdx5 | PMID: 29497036 | Prx5 transcript is highly expressed early along the transition of hematopoietic stem cells to mature neutrophils |
| S100a8 | PMID: 28137827 | Works alongside S100a9, involved in inflammatory responses in myeloid cells. S100A8/A9 induces neutrophil activation, including adhesion, CD11b upregulation, and shedding of CD62L. |
| C1qbp | PMID: 34860557 | Associated with pre-mRNA splicing factor and can promote differentiation of immune cells via metabolic-epigenetic reprogramming |
| Cxcr4 | PMID: 29734477<br>PMID: 29466759 | Cxcr4 is a master regulator of neutrophil development, proliferation and retention. |

*Supplementary Table 1. Significant genes identified during the critical period of cell decision-making (highlighted as the yellow box in Figure 3). The table includes columns for gene names, corresponding references to previous studies, and a description of their roles.*
